# Mining Archive.org’s Twitter Stream Grab for Pharmacovigilance Research Gold

**DOI:** 10.1101/859611

**Authors:** Ramya Tekumalla, Javad Rafiei Asl, Juan M. Banda

## Abstract

In the last few years Twitter has become an important resource for the identification of Adverse Drug Reactions (ADRs), monitoring flu trends, and other pharmacovigilance and general research applications. Most researchers spend their time crawling Twitter, buying expensive pre-mined datasets, or tediously and slowly building datasets using the limited Twitter API. However, there are a large number of datasets that are publicly available to researchers which are underutilized or unused. In this work, we demonstrate how we mined over 9.4 billion Tweets from archive.org’s Twitter stream grab using a drug-term dictionary and plenty of computing power. Knowing that not everything that shines is gold, we used pre-existing drug-related datasets to build machine learning models to filter our findings for relevance. In this work we present our methodology and the 3,346,758 identified tweets for public use in future research.

## Introduction

The World Health Organization (WHO) defined Pharmacovigilance as “the science and activities relating to the detection, assessment, understanding and prevention of adverse effects or any other drug-related problem” ^1^. The aim of pharmacovigilance is to enhance patient care and safety in relation to the use of medicines; and to support public health programmes by providing reliable, balanced information for the effective assessment of the risk-benefit profile of medicines. Traditionally, clinical trials are employed to identify and assess the profile of medicines. However, since they have limited ability to detect all ADRs due to factors such as small sample sizes, relatively short duration, and the lack of diversity among study participants, post marketing surveillance is required ^2^. The Food and Drug Administration (FDA) provides several post marketing surveillance programs like FDA Adverse Event Reporting System (FAERS), MedWatch to report events, however, under-reporting limits its effectiveness. A review of 37 studies established that more than 90% of ADRs are estimated to be under-reported ^3^. Social media platforms like Twitter and Facebook contain an abundance of text data that can be utilized for pharmacovigilance ^4^. Many studies presented satisfactory results by utilizing social media for pharmacovigilance and helped create a curated dataset for drug safety ^5^. Several other studies presented a connection between drugs and addictive behaviour among students using Twitter ^6,7^. However, it is challenging to use Twitter due to limitations with service providers who can export only 50,000 tweets per day. Further, usage of Twitter’s API or software to extract tweets is extremely time-consuming and economically unviable, especially for obtaining tweets relevant to a particular domain. Additionally, machine learning and deep learning models need exorbitant amounts of training data to train a model and not much data is available publicly for training.

In this context, data sharing ^8^ is a novel idea for research parasites to scavenge available datasets and apply their methodologies. Recent studies ^9–11^ prove that data sharing improves quality and strengthens research. The increase in collaborative efforts will provide an opportunity for researchers to continually enhance research ideas and avoid redundant research efforts ^12,13^. As part of this research, we scavenged a large publicly available dataset and procured data related to pharmacovigilance. In this paper, we present a data corpus of **3,346,758** carefully filtered tweets. The deliverables ^14^ include filtered tweet ids list, code to download and separate tweets from the Internet Archive (IA). Due to Twitter’s terms of service, tweet text cannot be shared. Therefore, only tweet ids are publicly made available. The whole methodology can be reproduced using the deliverables. This corpus can help train machine learning and deep learning models and can be reused by other researchers to enhance research in Pharmacovigilance.

### Data Preparation

The Internet Archive (IA) ^15^ is a non-profit organization that builds digital libraries of Internet sites and other cultural artifacts in digital form and provides free access to researchers, historians and scholars. For this research, we used the largest publicly available Twitter dataset in Internet Archive, which contains several json files of tweets in tar files sorted by date for each month of the year. The tar file must be downloaded and decompressed before usage. A total of 9,406,233,418 (9.4 billion) tweets for the years 2012 to 2018 are available in this dataset, we filtered this data using a drug terms dictionary to identify drug-specific tweets. The time taken to download, process and filter these tweets was 132 days.

### Drug Dictionary Creation

The UMLS^16^ is a large, multi-purpose and multilingual vocabulary database that contains information about biomedical and health related concepts, their various names, and the relationships among them. The UMLS includes the Metathesaurus, the Semantic Network, and the SPECIALIST Lexicon and Lexical Tools. Metathesaurus, the biggest component of UMLS was utilized in creating the drug dictionary, more specifically the RxNorm^17^ vocabulary. This vocabulary provides normalized names for clinical drugs and link names to the drug vocabulary commonly used in pharmacy management and drug interaction software. The MRCONSO table was filtered using RxNorm and Language of Term (LAT), which was set to English. The filtered table contained a total of 279,288 rows. Since, the dictionary was used on Twitter data and the total number of characters allowed in a tweet was 140 (until 2017) and 280 (from October 2017 onwards), we eliminated all the strings of length less than or equal to 3 (too ambiguous) and greater than or equal to 100. This was due to a less likely chance for tweets to contain drug names that were as short as 3 characters or as long as 100 characters. Further, we removed strings such as “2,10,15,19,23-pentahydrosqualene” which are chemical compounds. This elimination was based on the premise that users would find it cumbersome and tedious to type detailed chemical names of drugs, especially on social media. Additionally, we removed 50 terms like “disk, foam, bar-soap, sledgehammer, cement, copper, sonata” as these terms are not commonly used as drug names and in pharmacovigilance. After deleting the common terms and chemical compounds, only 266,556 rows were available of which five term types were used in the drug dictionary for the research. The dictionary also consists of a Concept Unique Identifier (CUI) to which strings with the same meaning are linked. The CUI is used in order to ensure that the meanings are preserved over time regardless of the different terms that are used to express those meanings. All the strings have been converted to lowercase and trimmed of white spaces. A total of 111,518 unique strings were used in total to create the drug dictionary. Table 1 represents the number of strings used for each term type and Table 2 contains sample rows from drug dictionary.

**Table 1:** Term types, their definitions and Number of strings

| Term Type | Example | # Strings |
| --- | --- | --- |
| Ingredients (IN) | Fluoxetine | 11,427 |
| Semantic Clinical Drug Component (SCDC) | Fluoxetine 4 MG/ML | 27,038 |
| Semantic Branded Drug Component (SBDC) | Fluoxetine 4 MG/ML [Prozac] | 17,938 |
| Semantic Clinical Drug (SCD) | Fluoxetine 4 MG/ML Oral Solution | 35,112 |
| Semantic Branded Drug (SBD) | Fluoxetine 4 MG/ML Oral Solution [Prozac] | 20,003 |

**Table 2:** Sample Drug Dictionary

| Concept Unique Identifier (CUI) | Term String |
| --- | --- |
| C0290795 | adderall |
| C0700899 | benadryl |
| C0025219 | melatonin |
| C0162373 | prozac |
| C0699142 | tylenol |

## Methods

In order to identify drug-specific tweets that would be useful for pharmacovigilance, we applied the drug dictionary on the Internet Archive twitter dataset. We filtered the dataset using spaCy, an open-source library in Python. We used the matcher in spaCy which would match sequences of tokens, based on pattern rules. Subsequently, the program generates an output file with the filtered tweets if it finds a match with the drug dictionary in the tweet text. Tweets are retrieved only if their language is set to English and if they are not retweeted. Initially, the method was performed on 2018 data, with our results showing that the maximum number of tweets that got separated consisted of a single drug string (one term). We speculate that, since Twitter has a limitation on the number of characters, people tend to write abbreviated terms or single terms that are either drug names or ingredients, instead of a drug string that consists of 4 or 5 terms. A single string dictionary is created from the five dictionaries with a total of 13,226 unique single terms. A total of 6 programs are run on each month for the year 2018. Only 10 months of data was available for the year 2018. Number of tweets obtained for four months for the year 2018 when used on six dictionaries are presented in table 3.

**Table 3:** Number of tweets obtained for each month from the Internet Archive Dataset in 2018.

| Month | Number of Tweets available | Single string tweets | Ingredients tweets | SCD tweets | SCDC tweets | SBD tweets | SBDC tweets |
| --- | --- | --- | --- | --- | --- | --- | --- |
| January | 134,747,413 | 83,583 | 27,718 | 0 | 15 | 0 | 0 |
| August | 141,132,076 | 67,227 | 21,335 | 0 | 13 | 0 | 0 |
| September | 133,068,824 | 64,123 | 22,230 | 0 | 22 | 0 | 0 |
| October | 132,297,280 | 68,221 | 22,955 | 0 | 17 | 0 | 0 |
| **Total** | **1,102,507,263** | **385,503** | **196,788** | **1** | **112** | **0** | **0** |

The SCD, SBD and SBDC dictionaries did not yield any tweets from 2018. In order to determine the reason, we examined and analyzed the dictionaries. For each term type in the drug dictionary, we calculated the lengths of all drug strings and identified the number of characters at each length ranging between 4 to 99 characters. Further, we also noted the average and median lengths of the drug strings. Table 4 depicts detailed statistics for each term type.

**Table 4:**
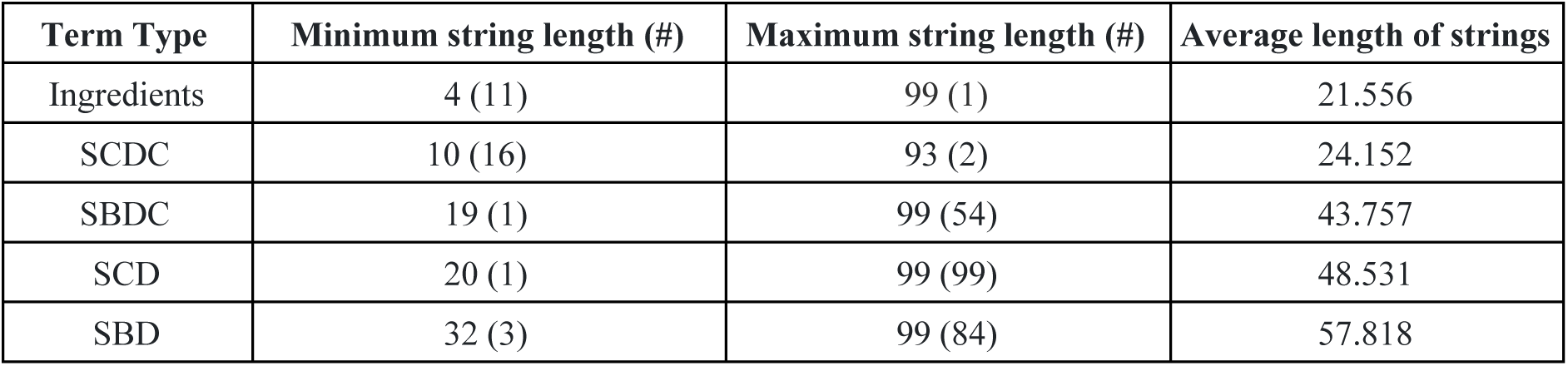
Statistics for each drug term type.

| Term Type | Minimum string length (#) | Maximum string length (#) | Average length of strings |
| --- | --- | --- | --- |
| Ingredients | 4 (11) | 99 (1) | 21.556 |
| SCDC | 10 (16) | 93 (2) | 24.152 |
| SBDC | 19 (1) | 99 (54) | 43.757 |
| SCD | 20 (1) | 99 (99) | 48.531 |
| SBD | 32 (3) | 99 (84) | 57.818 |

SBD, SCD and SBDC had the highest number of lengthy drug strings. The following drug string from SCDC drug dictionary, “***pneumococcal capsular polysaccharide type 33f vaccine 0.05 mg ml***”, has 64 characters. It is impractical to type the whole drug string in a tweet without an error. 90% of the tweets obtained from the SCDC were advertisements on either promoting the product or selling the product. Further examining all the tweets, we eliminated the 4 dictionaries (SCD, SCDC, SBD, SBDC) and used the single string and ingredients dictionary since it saves an enormous amount of computation time.

The following table represents the number of tweets retrieved for the years from 2012 to 2018 when used with the two remaining drug dictionaries.

**Table 5:** Number of single and ingredient tweets obtained from total tweets

| Month | Total tweets | Single string | Ingredients |
| --- | --- | --- | --- |
| 2018 | 1,102,507,263 | 588,854 | 26,034 |
| 2017 | 1,448,114,354 | 1,079,616 | 52,058 |
| 2016 | 1,427,468,805 | 1,252,057 | 62,514 |
| 2015 | 1,224,040,556 | 1,149,736 | 55,367 |
| 2014 | 1,086,859,898 | 873,937 | 47,953 |
| 2013 | 1,871,457,526 | 1,463,184 | 85,618 |
| 2012 | 1,245,785,016 | 76,795 | 4,139 |
| **Total tweets** | **9,406,233,418** | **6,152,862** | **314,680** |

We make the code publicly available for reproducibility. A total of 132 days were required to download, unzip and filter the tweets using drug dictionary. For all the years, each month was downloaded individually, unzipped to retrieve the json file and then the tweets were filtered using the drug dictionary. Typically, for a month, the method would require 10 minutes to download, 5 hours to unzip and 2 days to filter tweets on an IBM Blade Server with 768 GB RAM, 2 x Intel® Xeon® E5-269880 Processors, with 40 cores each, and 12TB of hard disk space.

The single string and ingredients dictionary was used on the IA dataset and a total of 6,703,331 (6.7 million) tweets were retrieved from 9,406,233,418 (9.4 billion) tweets. After eliminating duplicate tweets, a total of **6,703,166** were retrieved. We examined the retrieved tweets and found that more than 50% of the tweets are not relevant to pharmacovigilance. This is because some drug strings are used in common terminology and in other fields like math, technology etc. For example, the drug string “tablet” was used in reference to the electronic gadgets (Samsung, Microsoft tablets). In order to eliminate the tweets that are relevant to other domains and not pharmacovigilance, we employed machine learning and deep learning classification models to filter tweets.

## Classification

Since the filtered tweets contain a number of irrelevant tweets, we experimented with several classical machine learning and deep learning models on the filtered tweets to clean the tweets. The tweets obtained after classification can be used for training different machine learning and deep learning models by other researchers. Since there are no trainable datasets that we could make use of, we created a dataset utilizing annotated datasets from different sources ^18–21^. We emphasize that we did not annotate or create any annotated set of tweets ourselves.

### Classical Models

We collected 259,042 tweets that only have drug strings from multiple papers on pharmacovigilance using social media ^18–21^ and downloaded all the tweets available through them. These tweets were annotated by different annotators as part of their research. The collected 259,042 tweets from multiple pharmacovigilance papers were labelled as “drug” tweets. Additionally, we randomly collected 300,208 non-drug tweets from multiple years from Internet Archive and labelled them as “non-drug” tweets. Pre-processing was performed on the downloaded tweets by removing links and emojis and only tweet text was separated. A total of 559,250 tweets were used as an annotated training set, where only drug tweets were the actual annotated tweets collected from different sources. We experimented with five classifiers: Naive Bayes, Logistic Regression, Support Vector Machines (SVM), Random Forest and Decision Trees using the scikit-learn^22^. Support-Vector Machine constructs a hyperplane or set of hyperplanes in a high- or infinite-dimensional space, which can be used for classification, regression, or other tasks like outliers detection. We used a LinearSVC which is similar to SVC, but implemented in terms of liblinear rather than libsvm, so it has more flexibility in the choice of penalties and loss functions and should scale better to large numbers of samples. Naive Bayes methods are a set of supervised learning algorithms based on applying Bayes’ theorem with the “naive” assumption of conditional independence between every pair of features given the value of the class variable. We used the Multinomial Naive Bayes which implements the naive Bayes algorithm for multinomial distributed data and is one of the two classic naive Bayes variants used in text classification. A Random Forest is a meta estimator that fits a number of decision tree classifiers on various sub-samples of the dataset and uses averaging to improve the predictive accuracy and control over-fitting. The DecisionTreeClassifier uses a CART algorithm (Classification And Regression Tree). CART is a non-parametric decision tree learning technique that produces either classification or regression trees, depending on whether the dependent variable is categorical or numeric, respectively. However, the scikit uses an optimized version of the CART which does not support categorical values.

Each classifier model is applied on the stratified 75-25% (training - test) split of the annotated training set. We calculated precision, recall and F-score to evaluate each classifier and the results are tabulated in Table 6.

**Table 6:**
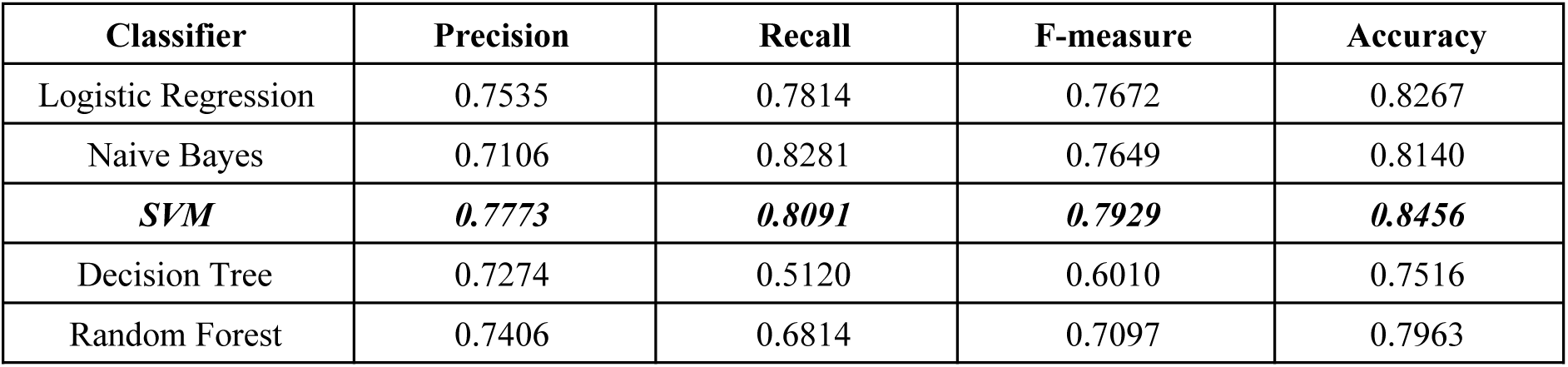
Classification metrics for the classical machine learning models

### Deep Learning models

For classification task, we experimented with six deep learning techniques from the Matchzoo framework ^23^: MVLSTM, DUET, KNRM, CONVKNRM, DSSM, and ARC-II ^24–29^. DUET is applied for the document ranking task and is composed of two separate deep neural networks, one matches the query and the document using a local representation, and another matches the query and the document using learned distributed representations. KNRM is a kernel-based neural model for conducting the document ranking task by using three sequential steps: 1) a translation matrix to model word-level similarities using word embeddings. 2) a modern kernel-pooling technique to use kernels for multi-level soft match features extraction. 3) a learning-to-rank layer that combines those features into the final ranking score. CONVKNRM uses CNNs to compose n-gram embeddings, and cross-matches n-grams of various lengths. It applies kernel pooling to extract ranking features, and uses learning-to-rank to obtain the final ranking score. MVLSTM conducts sentence matching with multiple positional sentence representations where each representation is generated by a bidirectional LSTM. The final score is produced by aggregating interactions between these different positional sentence representations. ARC-II focuses on sentence matching by naturally combining the hierarchical sentence modeling through layer-by-layer composition and pooling and capturing of the rich matching patterns at different levels of abstraction. DSSM aims to rank a set of documents for a given query. First, a non-linear projection maps the query and documents to a common semantic space. Then, the relevance of a document with the query is calculated as cosine similarity between their vectors in the semantic space. These deep models are general-purpose models and can be used for different text matching tasks such as document retrieval, conversational response ranking, and paraphrase identification. Precision, recall, F-measure, and accuracy metrics are tabulated in Table 7.

**Table 7:** Classification metrics for the deep learning models

| **Classifier** | **Precision** | **Recall** | **F-measure** | **Accuracy** |
| --- | --- | --- | --- | --- |
| **ARC-II** | **0.9679** | **0.9581** | **0.9630** | **0.9731** |
| DUET | 0.9635 | 0.9579 | 0.9607 | 0.9715 |
| DSSM | 0.6457 | 0.3904 | 0.4866 | 0.7001 |
| KNRM | 0.9585 | 0.9158 | 0.9367 | 0.9549 |
| MVLSTM | 0.9781 | 0.9047 | 0.9400 | 0.9579 |
| CONVKNRM | 0.9692 | 0.9402 | 0.9545 | 0.9673 |

### Calculating cutoff thresholds

We applied all classical and deep learning models on the filtered 6 million tweets to predict the probability score of each tweet from Internet Archive dataset. In order to determine what is the most optimal probability cutoff, we applied mixture models concepts ^30^. The way this methodology works is by taking all the probability scores and divide them into several hundreds of bins. A histogram of probability frequency is determined by calculating the number of observations in each bin. Based on hypothesis of mixture models, probability scores are distributed according to a mixture of two Gaussian distributions (drug and non drug tweets). Finally, two highest peaks of two Gaussian distributions and one valley with most depth between the two peaks are detected and the valley’s deepest point is used as cut-off point (threshold). In Figure 1 and 2, we plot the number of tweets that have a given probability score. Starting from an assigned probability of one, we cumulatively count the number of tweets we would keep at any given probability threshold. These plots allow us to see the selectivity of each model and the number of tweets at each threshold limit. Note that the optimal cutoff threshold is displayed next to the model name in the figure legend.

**Figure 1.**
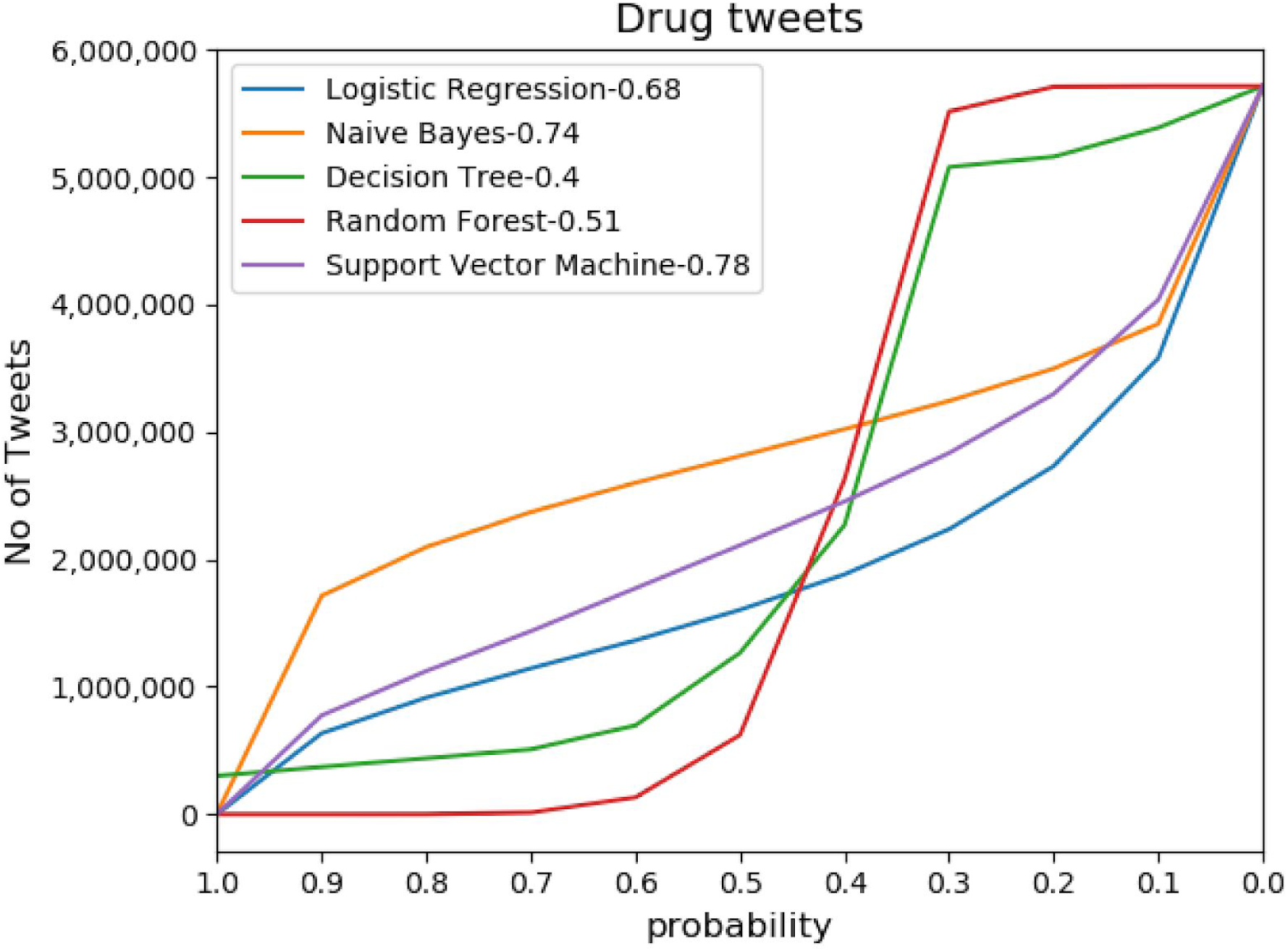
Probabilities of drug tweets using ML models.

**Figure 2.**
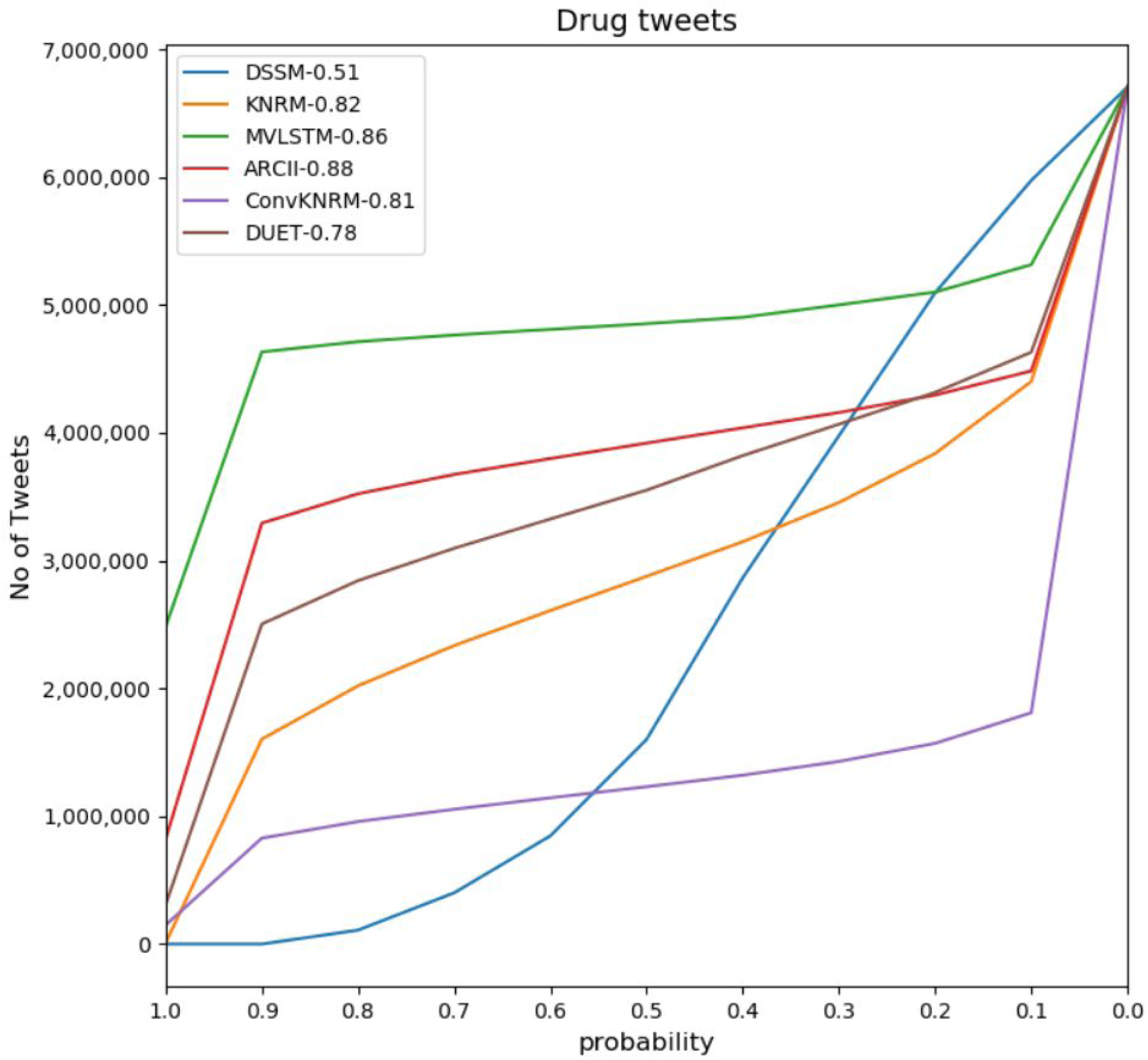
Probabilities of drug tweets using deep learning models.

Based on Tables 6 and 7, and the cutoff Figures 1 and 2, we selected **ARC-II** as the model to use to classify the relevance of the tweets. This deep learning model performed the best in terms of F-measure and accuracy, two of the metrics we deemed most relevant to identify useful tweets. After the classification filtering, we examined all the retrieved tweets and calculated the drug occurrences. We identified 6,867 unique drug strings in 3,346,758 million tweets. The entire methodology of the research is depicted in Figure 3 and Figure 4 depicts the popular drug strings and the number of occurrences for each drug string in the classified tweets.

**Figure 3:**
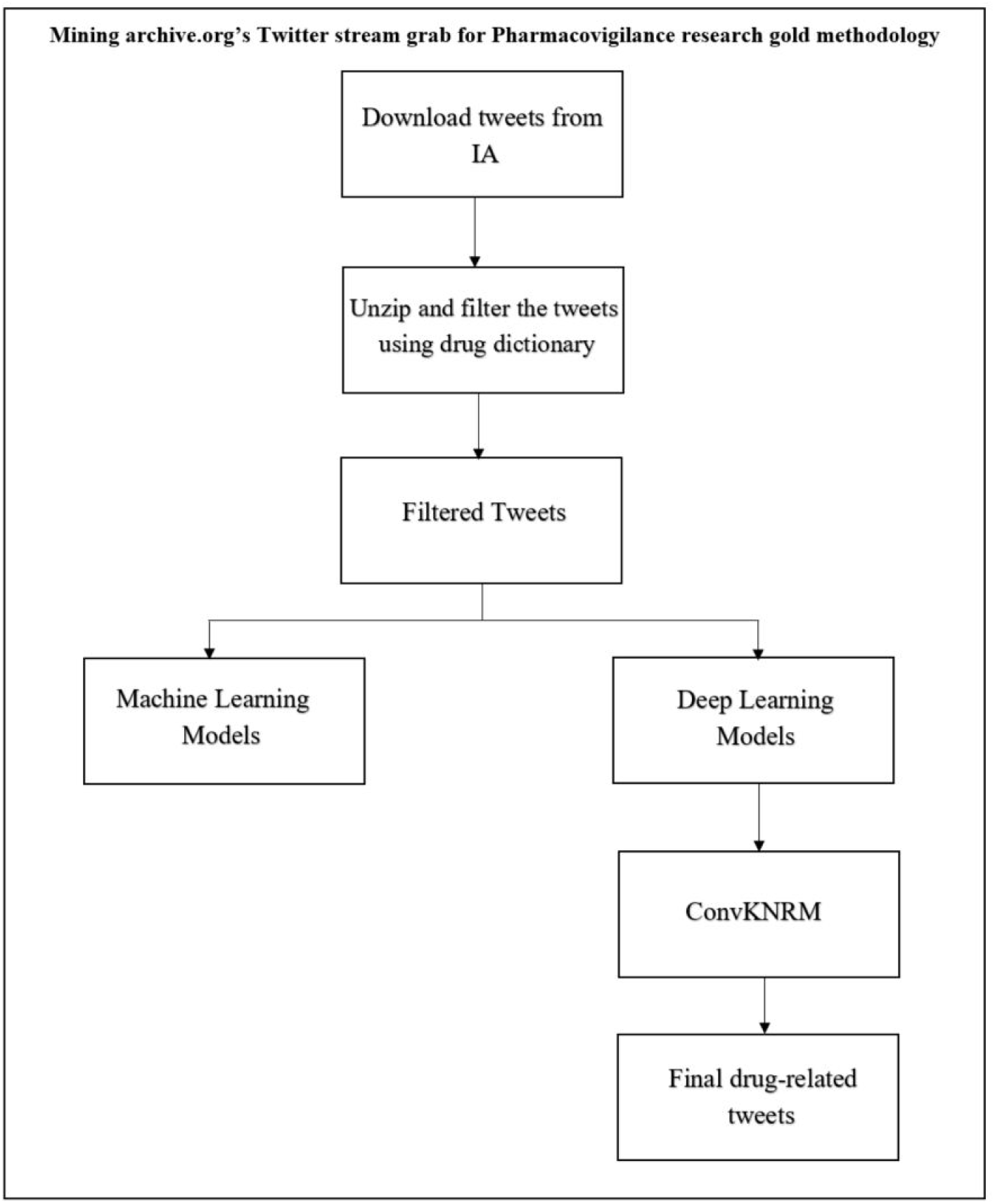
Methodology of Mining archive.org’s Twitter stream grab for Pharmacovigilance research gold

**Figure 4.**
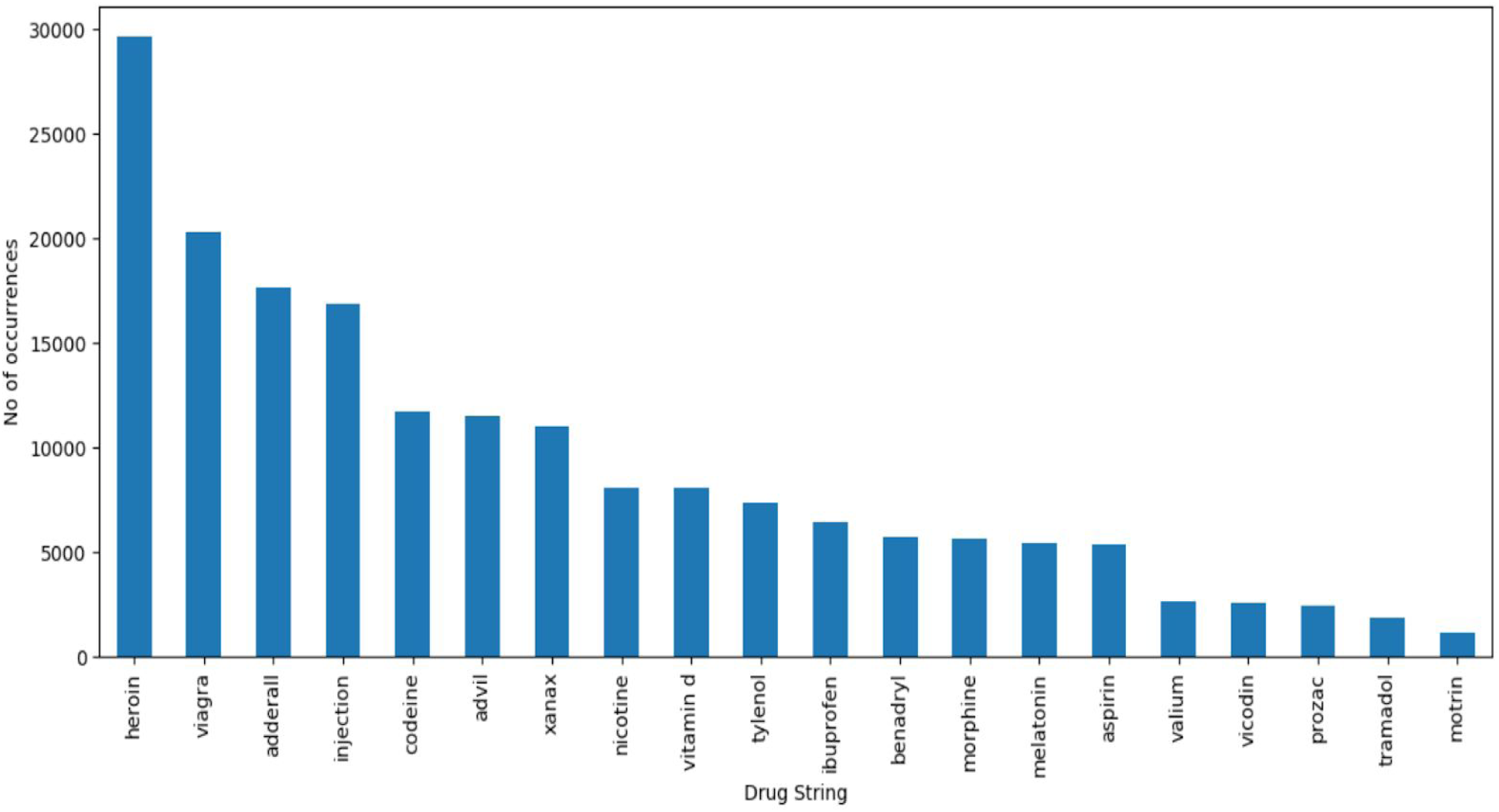
Popular drug string occurrences after filtering and classification of tweets.

Cocaine was the most popular drug string with 59,397 occurrences, however we eliminated it from the plot since it was used as a recreational drug than a medical drug. The following are a few examples of the tweets. The drug strings are highlighted in bold.

1. “my head still hurting tho. my mama gave me some **ibuprofen**. ”
2. “im so stupid for taking **benadryl** in the morning #sleeeeeepy”
3. “having to stop the **vicodin** completely because its making me sick seriously anything. #letthepainbegin”.
4. “i got very sick from **effexor**. also lost 2 years but ive recovered fully and life’s better than ever. hang in there”
5. “i took some **tylenol** # 3 with **codeine**. im sleepy, but i have to change the gauze in my mouth because it wont stop bleeding.”

### Future Work

The proposition of this paper is to utilize publicly available resources and employ Machine and Deep Learning techniques to create a dataset that can be made available for pharmacovigilance research. We believe that we can’t keep training models with very limited amount of manually annotated tweets, but we can use the theory of noisy labeling to create more robust models with silver standards ^31–33^. However, there are a few limitations which we would like to address in our future work. This research utilizes only English tweets since there were no publicly available annotated drug tweets for other languages. Currently, validation is performed only on the classification model but not the annotated dataset. Furthermore, the annotated drug tweets used in the training data were collected from publicly available sources and are labelled as drug tweets. Hence, edge cases such as ambiguous tweets were not considered. In the future, we would like to develop an improved annotated dataset which can be utilized as a gold standard dataset, following a trimodal distribution of probabilities where the edge cases are considered.

### Conclusion

In this paper, we scavenged a publicly available Twitter dataset, Internet Archive, mining over 9.4 billion tweets. Using a simple drug dictionary and plenty of computing power, we filtered 6 million tweets with relevant drug terms in them. In order to determine the viability of the filtered tweets for research work, we used publicly available, manually and expertly curated tweet datasets to build classification models to identify the relevant (or similar) tweets in our dataset. Overall, these tasks took around 150 days for downloading, filtering and classification, in order to retrieve 3,346,758 tweets which can be used for drug safety research and as a training set for other supervised methods by researchers. Further, we believe that this approach can be reused and extended to several other domains by changing the dictionaries and the filtering mechanisms.

